# APE1 recruits ATRIP to ssDNA in an RPA-independent manner to promote the ATR DNA damage response

**DOI:** 10.1101/2022.08.12.503732

**Authors:** Yunfeng Lin, Jia Li, Haichao Zhao, Anne McMahon, Shan Yan

## Abstract

Cells have evolved the DNA damage response (DDR) pathways in response to DNA replication stress or DNA damage. In the ATR-Chk1 DDR pathway, it has been proposed that ATR is recruited to RPA-coated single-strand DNA (ssDNA) by direct ATRIP-RPA interaction. However, it remains elusive whether and how ATRIP is recruited to ssDNA in an RPA-independent manner. Here, we provide evidence that APE1 directly associates ssDNA to recruit ATRIP onto ssDNA in an RPA-independent fashion. The N-terminal motif within APE1 is required and sufficient for the APE1-ATRIP interaction *in vitro* and the distinct APE1-ATRIP interaction is required for ATRIP recruitment to ssDNA and the ATR-Chk1 DDR pathway activation in *Xenopus* egg extracts. In addition, APE1 directly associates with RPA70 and RPA32 via two distinct motifs. Taken together, our evidence identifies APE1 as a direct recruiter of ATRIP onto ssDNA independent of RPA in the activation of ATR DDR pathway.

**Summary:**

- APE1 associates with ssDNA and ATRIP directly via distinct motifs and thereby recruits ATRIP onto ssDNA independent of RPA to promote the ATR DDR.
- APE1 interacts with RPA via distinct two motifs *in vitro* but such RPA-APE1 interaction is dispensable for ATRIP recruitment onto ssDNA.

## Introduction

The DNA damage response (DDR) signaling pathways such as ATR-Chk1 and ATM-Chk2 are activated by DNA replication stress or different DNA damage to coordinate DNA repair with cell cycle as well as apoptosis and senescence (Bartek et al., 2004; Branzei and Foiani, 2010; Ciccia and Elledge, 2010; Cimprich and Cortez, 2008; Harper and Elledge, 2007; Harrison and Haber, 2006; Su, 2006). In response to stalled DNA replication forks and different DNA lesions including DNA double-strand breaks (DSBs) and single-strand breaks (SSBs), ATR DDR can be recruited to and activated by RPA-coated single-stranded DNA (ssDNA) derived from functional uncoupling of MCM helicase and DNA polymerase activities, DSB end resection in the 5’-3’ direction, or SSB end resection in the 3’-5’ direction (Ciccia and Elledge, 2010; Cimprich and Cortez, 2008; Lin et al., 2018; Marechal and Zou, 2015; Shiotani and Zou, 2009). ATR activation also requires several mediator proteins such as ATRIP, TopBP1 and the Rad9-Rad1-Hus1 (9-1-1) complex (Delacroix et al., 2007; Kumagai et al., 2006; Yan and Michael, 2009; Zou and Elledge, 2003). Activated ATR kinase then phosphorylates a variety of substrates such as Chk1, among others, and phosphorylated Chk1 is the activated version of Chk1 kinase to regulate cell cycle progression and often serves as an indicator of ATR DDR activation (Chen and Sanchez, 2004; Guo et al., 2000; Matsuoka et al., 2007).

Since the discovery of ATRIP in ATR DDR pathway about 20 years ago, it has been an active subject of studies regarding how the ATR-ATRIP complex is recruited to ssDNA and activated by ATR activator/mediator proteins to maintain genome integrity (Cortez et al., 2001; Zou and Elledge, 2003). Earlier studies from several groups using human and yeast cells as well as *Xenopus* egg extracts have revealed independently that ATR and ATRIP associate with each other into a tight complex and that the direct ATRIP recognition and interaction with RPA-ssDNA is essential for the recruitment of ATR to ssDNA regions at sites of DNA damage for ATR activation (Lee et al., 2003; Unsal-Kacmaz and Sancar, 2004; Zou and Elledge, 2003). However, the RPA requirement of ATRIP recruitment to ssDNA for ATR DDR activation is sort of questioned by several follow up reports demonstrating that ATR-ATRIP complexes can bind to ssDNA in an RPA-independent manner *in vitro*, and that the low affinity RPA-independent recruitment of ATRIP to ssDNA is mediated by an unknown protein in mammalian cell nuclear extracts (Bomgarden et al., 2004; Kim et al., 2005). Although the exact molecular determinants of ATRIP (such as the N-terminal 1-108 amino acid fragment of human ATRIP) interaction with RPA-ssDNA remain to be determined and reconciled (Ball et al., 2005; Namiki and Zou, 2006), additional lines of investigations have demonstrated that TopBP1 can directly activate the ATR-ATRIP complex in *Xenopus* egg extracts and reconstituted human proteins in a RPA-dependent and RPA-independent manner (Choi et al., 2010; Choi et al., 2007; Kumagai et al., 2006). Whereas a good progress has been made regarding the implication of post-translational modifications (PTMs) such as sumoylation and phosphorylation in ATRIP regulation in ATR DDR (Memisoglu et al., 2019; Wagner et al., 2019; Wu et al., 2014), it remains an outstanding question in the field of genome integrity of how exactly ATRIP is recruited to ssDNA in an RPA-dependent and/or -independent fashion for ATR DDR activation.

As the major AP endonuclease, AP endonuclease 1 (APE1) has fast AP endonuclease activity but slow 3’-5’ exonuclease and 3’-phosphodiesterase activities as well as 3’-5’ RNA phosphatase and exoribonuclease activities (Boiteux and Guillet, 2004; Burkovics et al., 2006; Chohan et al., 2015; Hadi et al., 2002; Tell et al., 2009; Wilson and Barsky, 2001). In addition to its role in redox regulation for transcription, APE1 plays essential roles in various DNA repair pathways (Li and Wilson, 2014; Tell et al., 2009). Whereas APE1-knockout mice are embryonic lethal, the underlying mechanism of APE1 in cell viability remains unclear (Fung and Demple, 2005; Masani et al., 2013; Xanthoudakis et al., 1996). Human APE1 is genetically altered and aberrantly expressed and localized in cancer patients and has become an emerging therapeutic target for various cancer therapy (Abbotts and Madhusudan, 2010; Al-Safi et al., 2012; Fishel and Kelley, 2007; Koukourakis et al., 2001; Sengupta et al., 2016; Thakur et al., 2014; Yoo et al., 2008). Recent pre-clinical and clinical studies have shown encouraging finding of APE1 inhibitor APX3330 in anti-cancer therapy in solid tumors (Caston et al., 2021; Shahda et al., 2019). Our recent studies have demonstrated that the ATR-Chk1 DDR pathway is activated by hydrogen peroxide-induced oxidative DNA damage and defined plasmid-based SSB structures in *Xenopus* High-Speed Supernatant (HSS) system (Lin et al., 2018; Wallace et al., 2017; Willis et al., 2013). To promote the ATR DDR activation, APE1 initiates the 3’-5’ end resection at SSB sites to generate a short ∼1-3nt ssDNA gap via its exonuclease activity, followed by PCNA-mediated APE2-dependent SSB end resection continuation (Lin et al., 2018; Lin et al., 2020). This APE1/2-mediated two-step processing of SSBs generates a longer stretch of ssDNA (∼18-26nt) coated by RPA, leading to the assembly of the ATR DDR complex (ATR-ATRIP, TopBP1, and 9-1-1 complex), subsequent ATR DDR activation, and eventual SSB repair (Hossain et al., 2018; Lin et al., 2018; Lin et al., 2020). However, it remains unknown whether APE1 plays a direct role in the ATRIP recruitment to ssDNA via a non-catalytic function in the presence and/or absence of RPA for the ATR DDR pathway.

Here, we provide direct evidence that in addition to its exonuclease-mediated function, APE1 plays a direct role in the recruitment of ATRIP to ssDNA in *Xenopus* egg extracts and in *in vitro* reconstitution system with purified proteins. The N-terminal motif of APE1 is required for its direct association with ssDNA *in vitro*, and such APE1-ssDNA interaction can be enhanced by RPA. Importantly, APE1 directly interacts with ATRIP and recruits ATRIP to ssDNA with the absence of RPA *in vitro*. A deletion mutant of APE1 deficient for ATRIP interaction but proficient for ssDNA association could not recruit ATRIP onto ssDNA in the presence of endogenous RPA in APE1-depleted *Xenopus* egg extracts. Similar to wild type APE1, a nuclease mutant APE1 still recruits ATRIP onto ssDNA in APE1-depleted HSS, suggesting that APE1’s role in ATRIP ssDNA recruitment is not dependent on its nuclease activity. APE1 directly associates with RPA *in vitro* via two distinct motifs within APE1. Notably, the RPA-interaction-deficient APE1 had no effect on the ATRIP recruitment onto ssDNA in *Xenopus* egg extracts. Overall, the data in this study demonstrating that APE1 is required for ATRIP recruitment to RPA-coated ssDNA for ATR DDR activation in *Xenopus* egg extracts, and that APE1 directly associates with and recruits ATRIP to ssDNA in the absence of RPA *in vitro*. These findings thus support a previously uncharacterized critical non-catalytic function of APE1 for direct ATRIP recruitment to ssDNA independent of RPA for the ATR DDR pathway.

## Results

### APE1 is required for the recruitment of ATRIP onto ssDNA in the ATR-Chk1 DDR pathway activation in *Xenopus* egg extracts

Our recent studies have revealed that APE1 plays an important role in the defined SSB structure-induced ATR-Chk1 DDR pathway via its 3’-5’ exonuclease activity in *Xenopus* HSS system (Lin et al., 2018; Lin et al., 2020). This APE1-mediated initiation of 3’-5’ SSB end resection (∼1-3nt ssDNA gap) will be followed by APE2 recruitment and activation to continue SSB end resection, generating a longer stretch of ssDNA (∼18-26nt ssDNA) coated by RPA for subsequent assembly of an ATR checkpoint protein complex including ATR-ATRIP, TopBP1, and 9-1-1 complex to activate ATR DDR (Lin et al., 2018; Wallace et al., 2017; Willis et al., 2013). In addition to the 3’-5’ SSB end resection initiation, we are interested in whether APE1 plays other roles such as non-catalytic function in the ATR DDR pathway. To test this question directly, we chose to utilize a defined plasmid DNA with a 1-3nt small ssDNA gap structure (designated as Gap plasmid) *in vitro* as previously described (Lin et al., 2018), and tested whether APE1 is still required for the ATR DDR in response to this Gap plasmid in HSS (Top of the panel of *Figure 1A*). As expected, the defined Gap plasmid (“Gap”), but not Control plasmid (“CTL”) triggered Chk1 phosphorylation, suggesting the activation of the ATR-Chk1 DDR pathway by the defined Gap plasmid in the *Xenopus* HSS (*Figure 1A*). Notably, the Gap plasmid-induced Chk1 phosphorylation was still impaired in APE1-depleted HSS system (“Extract”, *Figure 1A*), suggesting that APE1 may play an additional non-catalytic function in the ATR DDR pathway. To further define the additional role of APE1 in ATR DDR activation, we isolated the DNA-bound fractions from the HSS and examined the abundance of checkpoint proteins via immunoblotting analysis. Although the recruitment of RPA70 and RPA32 to the Gap plasmid was slightly reduced when APE1 was depleted from the HSS (“DNA-bound”, *Figure 1A*), the presence of RPA70 and RPA32 on Gap-plasmid DNA suggests that the gap structure can be further processed by APE2 in the APE1-depleted HSS, consistent with our previous studies (Lin et al., 2018; Lin et al., 2020). Notably, the recruitment of ATRIP onto Gap plasmid was significantly compromised in APE1-depleted HSS, suggesting that APE1 plays a more direct role in ATRIP recruitment to ssDNA regions in the defined Gap plasmid (*Figure 1A*).

**Figure 1.**
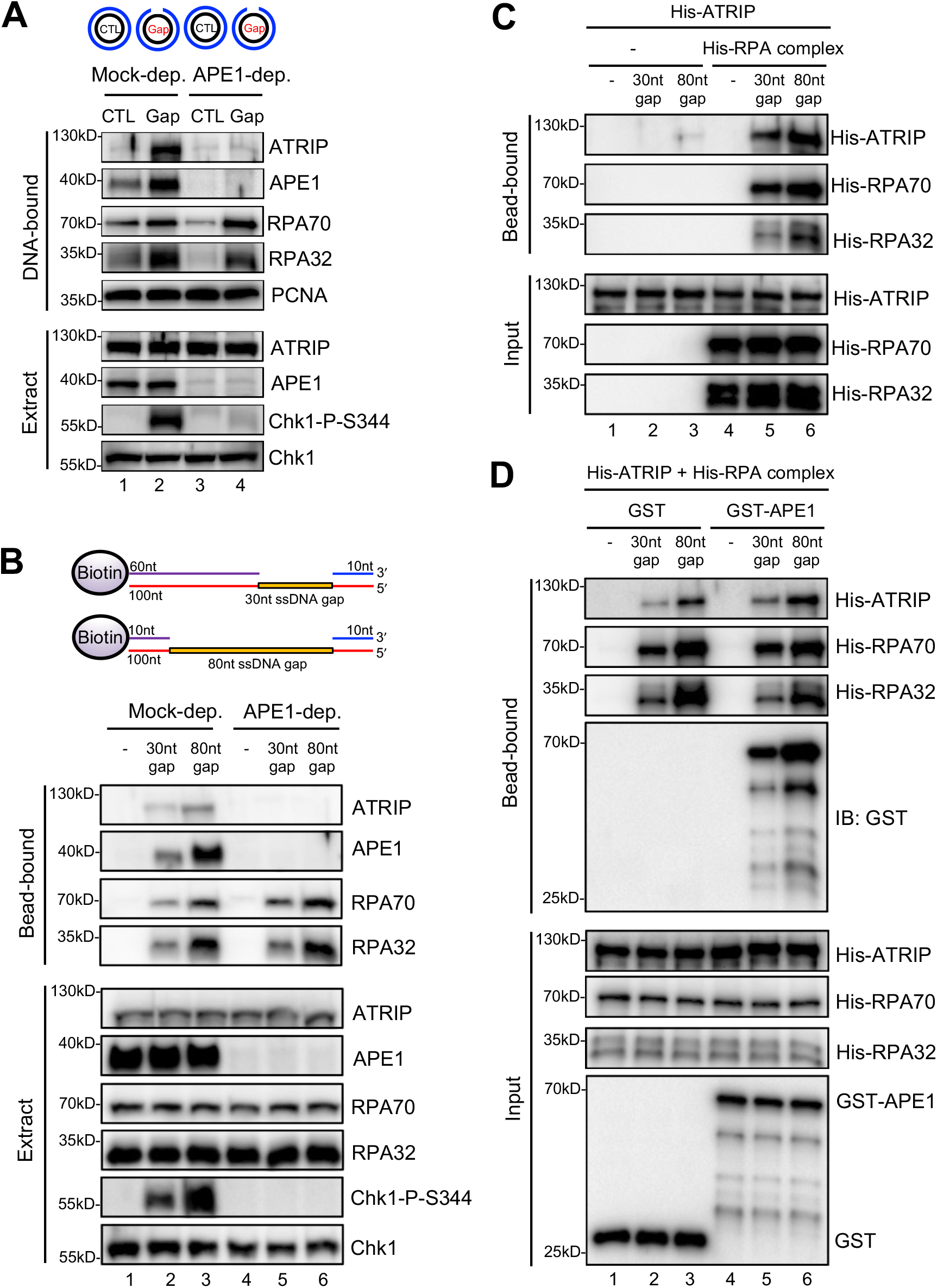
APE1 is required for the recruitment of ATRIP to RPA coated ssDNA in Xenopus egg extracts. (**A**) CTL (control) or Gap plasmid was added to Mock- or APE1-depleted HSS and incubated for 30 min. The DNA-bound fractions and total egg extract were examined via immunoblotting analysis as indicated. (**B**) Streptavidin beads coupled with equal moles of Biotin-labeled dsDNA with ssDNA gap structures (30nt or 80nt) were added to Mock- or APE1-depleted HSS. After incubation for 30 min at room temperature, the DNA-bound fractions and total egg extract were examined via immunoblotting analysis as indicated. (**C**) Streptavidin beads coupled with equal moles of Biotin-labeled dsDNA with ssDNA gap structures (30nt or 80nt) were added to an interaction buffer containing purified His-ATRIP protein with/without His-RPA protein. After incubation for 30 min at room temperature, the DNA-bound fractions and the input were examined via immunoblotting analysis. (**D**) Streptavidin beads coupled with equal moles of Biotin-labeled dsDNA with ssDNA gap structures (30nt or 80nt) were added to an interaction buffer containing His-ATRIP and His-RPA, which was supplemented with GST or GST-APE1. After incubation for 30 min at room temperature, the DNA-bound fractions and the input were examined via immunoblotting analysis.

In light of different lengths of ssDNA *in vitro* reconstitution systems by previous studies (such as 75nt or 80nt) (Bomgarden et al., 2004; Choi et al., 2010; Zou and Elledge, 2003), we intended to study the recruitment of ATRIP onto defined dsDNA structures with different length of ssDNA gaps. We chose to test two 100bp-dsDNA structures with either 30nt or 80nt ssDNA gap covalently linked with 5’-biotin on top strand (designated as “30nt gap” and “80nt gap”) for subsequent Streptavidin magnetic bead-bound isolation and analysis from *Xenopus* egg extracts (Top of the panel of *Figure 1B*). When equal moles of dsDNA with 30nt or 80nt ssDNA gap were added to HSS, more RPA70 and RPA32 as well as APE1 and ATRIP are recruited to the 80nt-ssDNA gap and Chk1 phosphorylation was also enhanced (*Figure 1B*). This enhanced Chk1 phosphorylation is likely due to increased RPA complex recruitment onto the 80nt-ssDNA gap (*Figure 1B*). Whereas APE1 depletion led to compromised Chk1 phosphorylation, the recruitment of ATRIP but not RPA70 nor RPA32 was compromised in APE1-depleted HSS (*Figure 1B*). Our observations so far suggest that APE1 is important for the recruitment of ATRIP onto RPA-coated ssDNA and Chk1 phosphorylation in *Xenopus* HSS.

Next, to determine whether APE1 plays any role in the RPA-dependent ATRIP recruitment onto ssDNA, we tested whether His-tagged ATRIP recombinant protein can be recruited to 30nt or 80nt ssDNA gap *in vitro* in the absence or presence of equal moles of recombinant RPA complex. Consistent with previously reported RPA-dependent ATRIP recruitment to ssDNA (Zou and Elledge, 2003), 30/80nt-ssDNA coated with RPA70 and RPA32 significantly enhanced the recruitment of His-ATRIP *in vitro*, although almost no binding of ATRIP onto ssDNA (30nt and 80mt) was observed in the absence of recombinant RPA complex (*Figure 1C*). We noticed more binding of His-ATRIP onto 80nt ssDNA gap compared with 30nt ssDNA gap (*Figure 1C*). As expected, the recruitment of His-ATRIP onto 30nt and 80nt ssDNA was similar to each when same amount of ssDNA gap structures was coupled to beads (*Figure 1-figure supplement 1A*). Furthermore, the addition of GST-APE1 but not GST protein increased the recruitment of His-ATRIP onto ssDNA with the presence of His-RPA complex *in vitro* (*Figure 1D*). It is worth to note that the presence of GST-APE1 had almost no noticeable effect on the recruitment of His-RPA70 and His-RPA32 to ssDNA gap structures (*Figure 1D*). Similarly, the presence of GST-APE1 but not GST increased the recruitment of endogenous ATRIP but not endogenous RPA70/RPA32 to ssDNA gap structures in the Xenopus HSS (*Figure 1-figure supplement 1B*). Whereas RPA itself is sufficient for ATRIP recruitment onto ssDNA *in vitro*, our observations here suggest that APE1 may stimulate the RPA-dependent ATRIP recruitment onto ssDNA *in vitro*. Alternatively, it is possible that APE1 may play an additional but direct role in the recruitment of ATRIP onto ssDNA *in vitro* that is independent of RPA.

### APE1 recognizes and binds with ssDNA directly in a length-dependent manner *in vitro*

Although APE1 is known as a DNA repair protein to specifically recognize and process AP site, it remains unclear whether and how APE1 interacts with ssDNA. To identify the possible direct role of APE1 in ATRIP recruitment onto ssDNA, we first performed systematic analysis of APE1 association with ssDNA. Our bead-bound experiments showed that GST-APE1 but not GST was recruited onto 30nt and 80nt ssDNA gap structures *in vitro* (*Figure 2A-B*). We also determined that GST-APE1 but not GST was recruited onto beads coupled with 70nt ssDNA *in vitro* (*Figure 2-figure supplement 1*). Furthermore, we demonstrated that GST-APE1 but not GST was recruited to beads coupled with 40nt, 60nt, and 80nt ssDNA, but not 10nt nor 20nt ssDNA (*Figure 2C*). Furthermore, the longer ssDNA is, the more GST-APE1 is recruited (*Figure 2C*). Collectively, these observations suggest an APE1-ssDNA interaction in a length-dependent manner *in vitro* (30-80nt) regardless the ssDNA is alone or in gapped structures.

**Figure 2.**
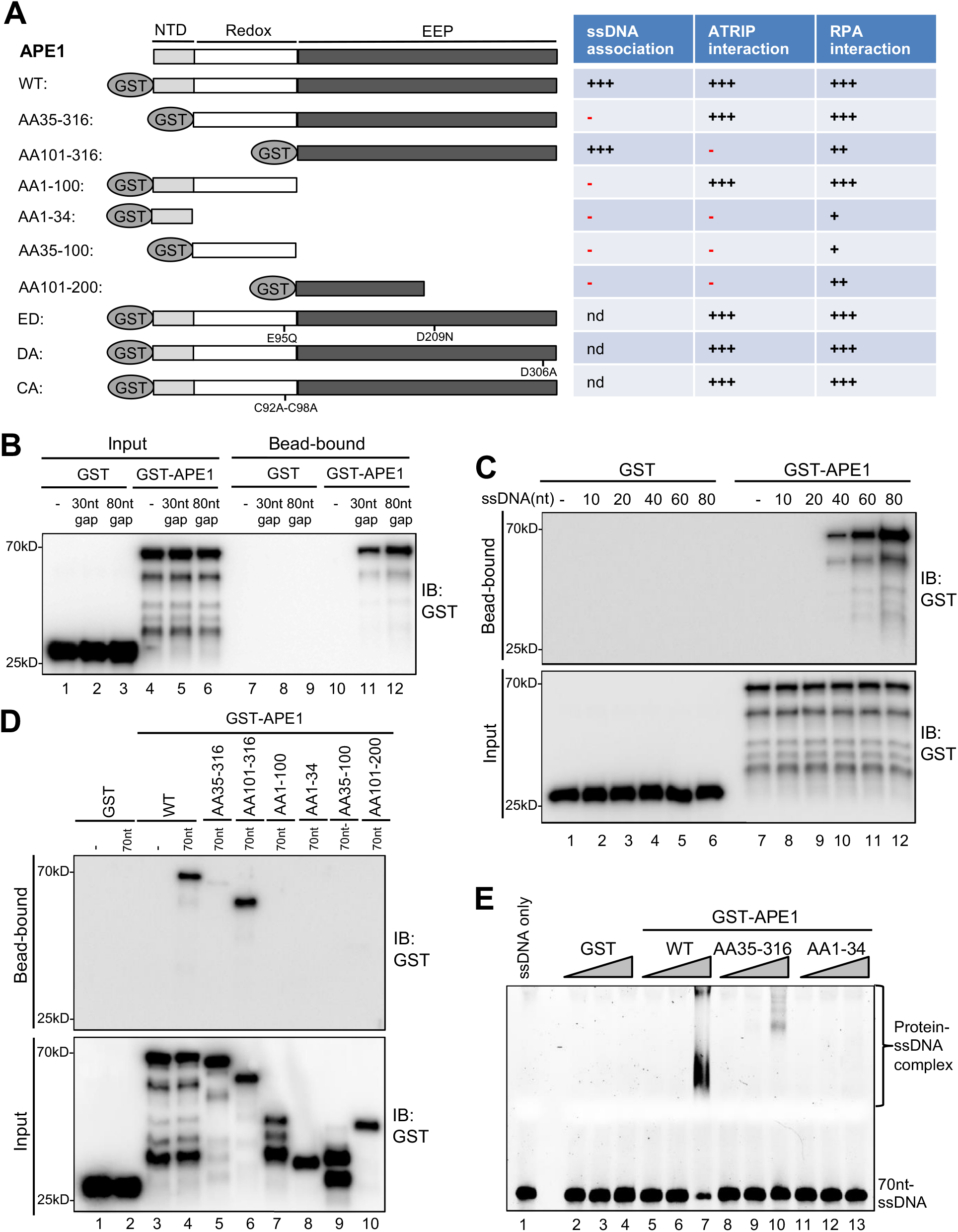
APE1 recognizes and binds with ssDNA in a length-dependent fashion *in vitro*. (**A**) Schematic diagram of APE1 functional domains and a summary of its interactions with ssDNA, ATRIP, and RPA from this study. (**B**) Streptavidin beads coupled with Biotin-labeled dsDNA with ssDNA gap structures (30nt or 80nt) were added to an interaction buffer containing GST or GST-APE1. After incubation for 30 min at room temperature, the DNA-bound fractions and the input were examined via immunoblotting analysis as indicated. (**C**) Streptavidin beads coupled with Biotin-labeled ssDNA with different lengths (10nt, 20nt, 40nt, 60nt, or 80nt) were added to an interaction buffer containing GST or GST-APE1. After incubation for 30 min at room temperature, the DNA-bound fractions and the input were examined via immunoblotting analysis. (**D**) Streptavidin beads coupled with Biotin-labeled ssDNA (70nt) were added to an interaction buffer containing (70nt) GST or WT or fragment of GST-APE1. After incubation for 30 min at room temperature, the DNA-bound fractions and the input were examined via immunoblotting analysis. (E) An EMSA assay shows the interaction between WT, AA35-316 and AA1-34 GST-APE1 and the 70nt-ssDNA structure *in vitro*.

To further dissect domain requirements within APE1 for ssDNA association, we generated a series of deletion GST-tagged APE1 and found that WT GST-APE1 and AA101-316 GST-APE1 but not any other deletion GST-APE1 tested (i.e., AA35-316, AA1-100, AA1-34, AA35-100, AA101-200) associated with beads coupled with 70nt-ssDNA *in vitro* (*Figure 2A and 2D*). Intriguingly, AA101-316 but not AA35-316 GST-APE1 associated with ssDNA (*Figure 2D*). We speculate that the fragment of AA35-100 within APE1 may somehow inhibit the APE1-ssDNA association due to a currently unknown mechanism. In addition, our EMSA assays revealed that WT GST-APE1 but not GST formed protein-ssDNA complex *in vitro* (*Figure 2E*). Notably, neither AA35-316 nor AA1-34 GST-APE1 formed protein-ssDNA complex in EMSA assays (*Figure 2E*). These observations suggest that AA1-34 within APE1 is required but seems not sufficient for ssDNA association at least under our tested conditions, and that APE1 AA35-316 is deficient for ssDNA association while APE1 AA101-316 is proficient in ssDNA interaction (*Figure 2*).

What are the effects of N-terminal motif of APE1 for its 3’-5’ exonuclease and AP endonuclease activities? Similar to our previous report (Lin et al., 2020), WT GST-APE1 but neither ED (E95Q-D306A) GST-APE1 nor GST displayed 3’-5’ exonuclease and AP endonuclease activities (*Figure 2-figure supplement 2A-B*). Notably, AA101-316 GST-APE1 is defective for 3’-5’ exonuclease and AP endonuclease activities (*Figure 2-figure supplement 2A-B*); however, AA35-316 GST-APE1 is proficient in AP endonuclease activity but deficient for 3’-5’ exonuclease activity (*Figure 2-figure supplement 2C-D*). These observations suggest the importance of the AA1-34 motif of APE1 for its 3’-5’ exonuclease activity and the AA35-100 motif within APE1 for its AP endonuclease activity.

### APE1 interacts and recruits ATRIP onto ssDNA in an RPA-independent manner *in vitro* and promotes the ATR DDR pathway in *Xenopus* egg extracts using a non-catalytic mechanism

We next tested whether and how APE1 might interact with ATRIP directly by protein–protein interaction assays. GST pulldown assays showed that GST-APE1 but not GST directly interacted with His-ATRIP *in vitro* (*Figures 2A and 3A*). Domain dissection experiments revealed that both AA35-316 GST-APE1 and AA1-100 GST-APE1 associated with His-ATRIP to the similar capacity as WT GST-APE1 (*Figure 3A*). However, AA101-316 GST-APE1 and other fragments of APE1 tested (i.e., AA1-34, AA35-100, and AA101-200) were deficient for interaction with His-ATRIP (*Figure 3A*). In addition, neither of the point mutants within GST-APE1’s active sites (i.e., ED, D306A, and C92A-C98A) affected the APE1-ATRIP interaction (*Figure 3B*), although they are deficient for 3’-5’ exonuclease as shown previously (Lin et al., 2020). Thus, our findings indicate that AA35-100 within APE1 is required for ATRIP interaction and AA1-100 is the minimum fragment within APE1 sufficient for ATRIP association *in vitro* (*Figures 2A and 3A*).

**Figure 3.**
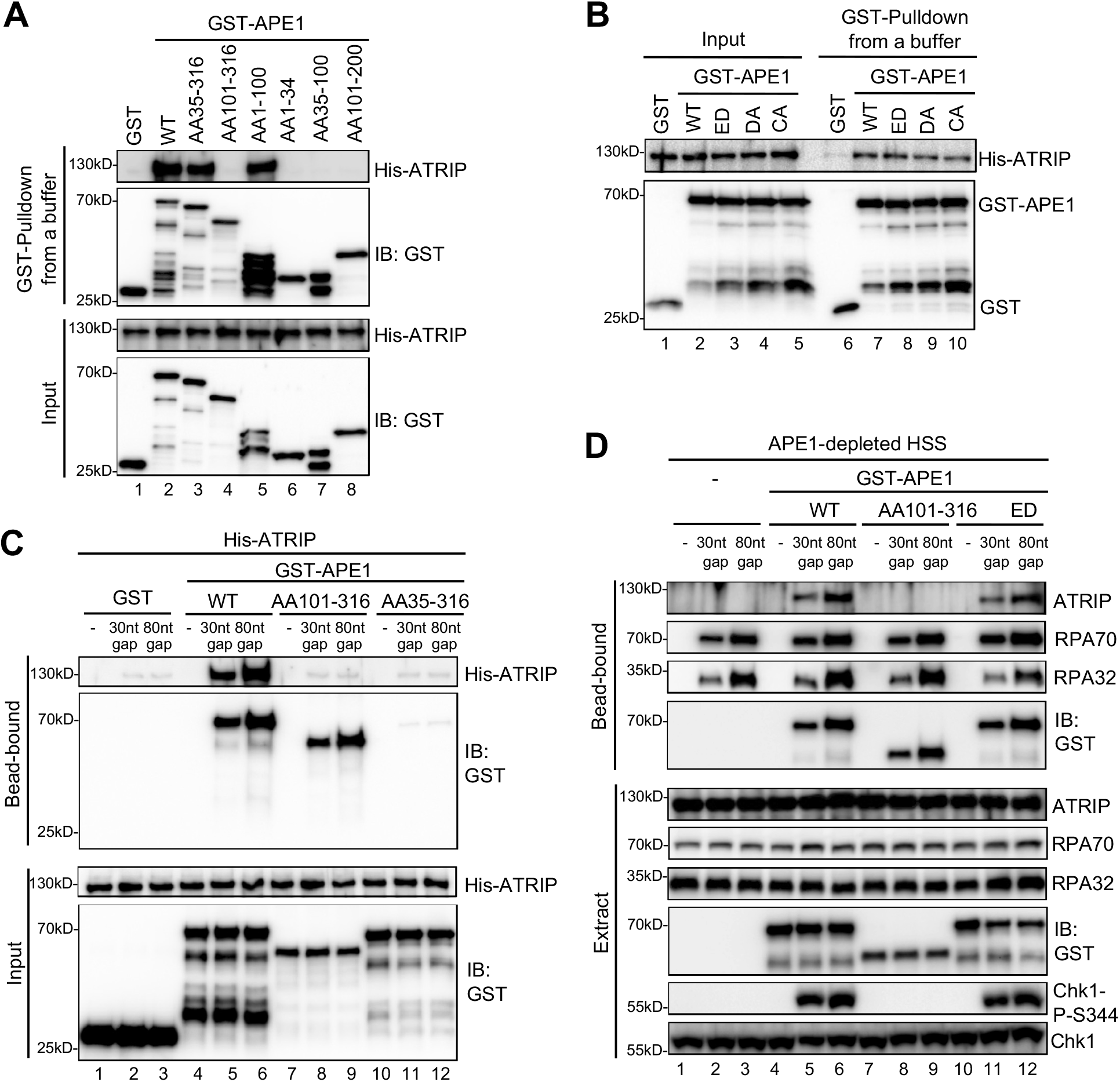
APE1 interacts and recruits ATRIP onto ssDNA in an RPA-independent manner *in vitro* and promotes the ATR DDR pathway in *Xenopus* egg extracts using a non-catalytic function mechanism. (**A-B**) GST pulldown assays with GST, WT or fragment/mutant GST-APE1 as well as His-ATRIP in an interaction buffer. The input and pulldown samples were examined via immunoblotting analysis. (**C**) Streptavidin beads coupled with Biotin-labeled dsDNA with ssDNA gap structures (30nt or 80nt) were added to an interaction buffer containing His-ATRIP and GST/GST-tagged proteins (WT, AA101-316, or AA35-316 GST-APE1) as indicated. DNA-bound fractions and Input samples were examined via immunoblotting analysis as indicated. (**D**) Streptavidin beads coupled with Biotin-labeled dsDNA with ssDNA gap structures (30nt or 80nt) were added to APE1-depleted HSS, which was supplemented with GST or GST-tagged proteins (WT, AA101-316, or ED GST-APE1) as indicated. DNA-bound fractions and total extract samples were examined via immunoblotting analysis as indicated.

Based on the observation of direct APE1-ATRIP interaction (*Figure 3A*), we intended to test whether APE1 could recruit ATRIP onto ssDNA directly in the absence of RPA *in vitro*. We found that His-ATRIP protein was recruited onto 30nt and 80nt ssDNA gap structures in the presence of WT GST-APE1 but not GST (Compare Lanes 4-6 and Lane 1-3 in “Bead-bound”, *Figure 3C*). Due to its deficiency in ssDNA interaction (*Figures 2D and 3A*), AA35-316 GST-APE1 was not recruited to 30nt and 80nt ssDNA gap structures, which led to the insufficient recruitment of His-ATRIP onto ssDNA (Lanes 10-12 in “Bead-bound”, *Figure 3C*). Notably, AA101-316 GST-APE1 was recruited to 30nt and 80nt ssDNA gap structures but could not recruit ATRIP to ssDNA, due to deficiency in ATRIP association (Lanes 7-9 in “Bead-bound”, *Figure 3C*). These observations strongly support that APE1 interacts with ssDNA via its AA1-34 fragment and recruits ATRIP onto ssDNA via its AA1-100 in *in vitro* reconstitution systems, and that such APE1-mediated ATRIP recruitment onto ssDNA is independent of RPA.

To test the biological significance of APE1-mediated ATRIP onto ssDNA, we performed rescue experiments in APE1-depleted HSS. WT GST-APE1 but not AA101-316 GST-APE1 rescued the recruitment of endogenous ATRIP onto 30nt and 80nt ssDNA gap structures and subsequent Chk1 phosphorylation, although endogenous RPA70 and RPA32 as well as WT/AA101-316 GST-APE1 associated with ssDNA gap structures in APE1-depleted HSS (Compare Lanes 4-6 and Lanes 7-9, *Figure 3D*). This observation indicates the significance of the APE1-ATRIP interaction for ATRIP recruitment onto ssDNA and subsequent ATR DDR pathway activation in *Xenopus* egg extracts.

In light of the significance of APE1 and its 3’-5’ exonuclease in the initial end resection of defined SSB structures and subsequent ATR DDR pathway in *Xenopus* HSS system (Lin et al., 2020), we sought to test whether APE1’s catalytic function plays a vital role in the direct ATRIP regulation and chose to use ED GST-APE1 lacking 3’-5’ exonuclease and AP endonuclease (*Figure 2-figure supplement 2A-B*) (Lin et al., 2020). We found that similar to WT GST-APE1, ED GST-APE1 bound with ssDNA and recruited endogenous ATRIP onto ssDNA and rescued Chk1 phosphorylation in APE1-depleted HSS (Lanes 10-12, *Figure 3D*). This observation suggests that APE1’s nuclease activity is dispensable for its direct recruitment of ATRIP onto ssDNA gap structures and subsequent ATR-Chk1 DDR pathway activation in the *Xenopus* HSS system. Together, these findings demonstrate that APE1 directly associates with and recruits ATRIP onto ssDNA *in vitro* and that APE1 recruits ATRIP onto ssDNA via a non-catalytic function to promote the ATR-Chk1 DDR pathway activation in the *Xenopus* HSS system.

### APE1 interacts with RPA70 and RPA32 via two distinct binding motifs

We have shown that both RPA complex and APE1 can independently recruit recombinant ATRIP onto ssDNA *in vitro* (*Figures 1C and 3C*), and that APE1 is required for the recruitment of ATRIP onto RPA-coated ssDNA in *Xenopus* HSS system (*Figure 1B*). Next, we sought to determine whether APE1 interacts with RPA directly, and, if so, whether RPA plays a role for APE1-mediated ATRIP recruitment onto ssDNA for ATR DDR in the HSS system. Due to the significant role of RPA in the recruitment of other DDR proteins such as TopBP1 to ssDNA (Acevedo et al., 2016), it is not feasible technically to test whether RPA depletion directly or indirectly affects APE1-mediated ATRIP recruitment and the subsequent ATR-Chk1 DDR pathway in the HSS system. Our strategy is to identify RPA-interaction motifs within APE1 *in vitro*, and to determine whether such a mutant APE1 deficient in RPA-interaction still recruits ATRIP onto ssDNA in the HSS system.

First, we tested the possibility that recombinant GST-APE1 might interact with purified recombinant His-RPA complex (RPA70, RPA32, and RPA14) by protein–protein interaction assays. Our GST pulldown assays showed that WT GST-APE1 but not GST interacted with His-RPA70 *in vitro* (*Figure 4A*). Almost no noticeable effects were observed for the interaction of ED, DA, and CA GST-APE1 with His-RPA70, suggesting that the E95, D209, D306, C92, and C98 residues in APE1 are not critical for ATRIP interaction (*Figure 4-figure supplement 1A*). Domain dissection experiments revealed that AA35-316 and AA1-100 GST-APE1 associated with His-RPA70 in a similar capacity to WT GST-APE1 (*Figure 4A*). The binding to His-RPA70 was decreased but nevertheless not completely eliminated in other deletion fragments of GST-APE1 tested (i.e., AA101-316, AA1-34, AA35-100, AA101-200) (*Figure 4A*). These observations suggest (I) that the first 100 amino acids of APE1 are important for RPA association and (II) that more than one binding sites within APE1 may mediate interaction with RPA complex.

**Figure 4.**
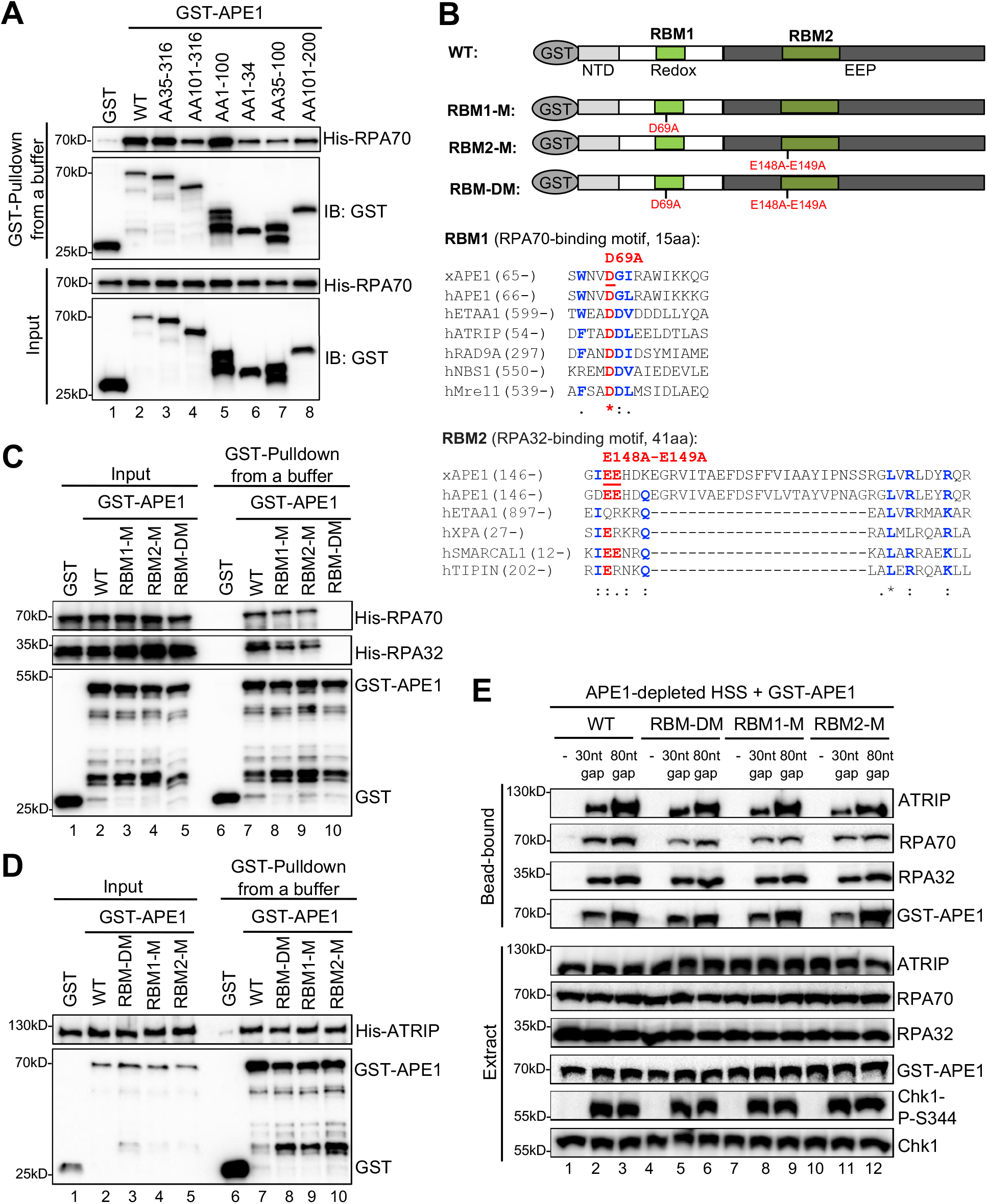
APE1 interacts with RPA70 and RPA32 via two distinct binding motifs. **(A)** GST pulldown assays with GST, WT or fragment of GST-APE1 as well as His-RPA in an interaction buffer. The input and pulldown samples were examined via immunoblotting analysis. **(B)** Schematic diagram of APE1 functional domains and its putative RPA binding motifs (RBM1 and RBM2), as well as sequence alignment of RBM1 and RBM2 from different RPA-interaction proteins. **(C)** GST pulldown assays with GST, WT/mutant GST-APE1 as well as His-RPA protein complex in an interaction buffer. The input and pulldown samples were examined via immunoblotting analysis. **(D)** GST pulldown assays with GST, WT/mutant GST-APE1 as well as His-ATRIP protein in an interaction buffer. The input and pulldown samples were examined via immunoblotting analysis. **(E)** Streptavidin beads coupled with Biotin-labeled dsDNA with ssDNA gap structures (30nt or 80nt) were added to APE1-depleted HSS, which was supplemented with WT or RBM mutant GST-APE1 (WT, RBM1-M, RBM2-M or RBM-DM GST-APE1) as indicated. DNA-bound fractions and total extract samples were examined via immunoblotting analysis as indicated.

Second, to further test our hypothesis of multiple bindings sites of APE1 for its interaction with RPA complex, we performed amino acid sequence alignments of APE1 (*Xenopus* APE1 and human APE1) to several human RPA70-interacting proteins (e.g., ETAA1, ATRIP, RAD9A, NBS1, and Mre11) and RPA32-interacting proteins (e.g., ETAA1, XPA, SMARCAL1, and TIPIN) and found that APE1 contains a putative RPA70-binding motif (15AA, designated as RBM1) and a putative RPA32-binding motif (41AA, designated as RBM2) (*Figure 4B*) (Bass et al., 2016; Haahr et al., 2016; Lee et al., 2016). To determine whether these two possible RPA-binding motifs within APE1 are important for RPA association, we generated single mutant in RBM1 (D69A, designated as RBM1-M) or RBM2 (E148A-E149A, designated as RBM2-M) or in combination (D69A-E148A-E149A, designated as RBM-DM) (*Figure 4B*). GST pulldown assays showed that RBM-DM GST-APE1 was defective for interaction with His-RPA70 and His-RPA32 *in vitro*, although RPA association was mildly impaired in single mutant RBM1-M and RBM2-M GST-APE1 (*Figure 4C*). However, neither of the RPA-binding mutants within APE1 affected its association with His-ATRIP protein (*Figure 4D*). In addition, neither of the RPA-binding mutant GST-APE1 (i.e., RBM1-M, RBM2-M, and RBM-DM) had noticeable effects on APE1’s 3’-5’ exonuclease and AP endonuclease activity, comparing with WT GST-APE1 under our experimetnal conditions (*Figure 4-figure supplement 2A-B*). Our observations here suggest that APE1 directly interacts with RPA70 and RPA32 *in vitro* using previously uncharacterized two distinct motifs within APE1 (i.e., RBM1 and RBM2) (*Figure 4B*).

Our earlier result showed that APE1-depletion led to defective ATRIP recruitment to ssDNA at gap structures and Chk1 phosphorylation in *Xenopus* HSS system (*Figure 1B*). Our rescue experiments showed that similar to WT GST-APE1, the RPA-interaction-deficient mutant GST-APE1 (RBM1-M, RBM2-M, and RBM-DM) rescued the recruitment of endogenous ATRIP and RPA70/RPA32 onto ssDNA and subsequent Chk1 phosphorylation in APE1-depleted HSS (*Figure 4E*). This result suggests that the RPA-APE1 interaction may be dispensable for the APE1-mediated ATRIP recruitment onto ssDNA in *Xenopus* egg extracts. In addition, we tested whether RPA plays any role in the APE1-ssDNA interaction in reconstitution system. Based on the length-dependent APE1 association with ssDNA (*Figure 2C*), we added excess recombinant His-RPA complex and found that APE1 interaction with longer ssDNA (40nt, 60nt, and 80nt) was enhanced by the presence of RPA complex *in vitro* (*Figure 4-figure supplement 1B*). Similar to WT GST-APE1, RBM-DM GST-APE1 was also recruited to longer ssDNA (40nt and 80nt) (*Figure 4-figure supplement 1C*); however, the RPA-stimulated ssDNA interaction of GST-APE1 was impaired when RBM-DM GST-APE1 was compared with WT GST-APE1 (*Figure 4-figure supplement 1C*). These observations suggest that RPA interaction may be important to stabilize the APE1-ssDNA interaction *in vitro*.

## Discussion

In addition to its critical roles in DNA repair and redox regulation (Li and Wilson, 2014; Tell et al., 2009), APE1 plays an essential role in the initiation step of 3’-5’ end resection in SSB-induced ATR-Chk1 DDR pathway via its 3’-5’ exonuclease acitivity (Lin et al., 2020). Mechanistic studies further elucidate that APE1 direclty recognizes and binds to SSB site and generate a small ssDNA gap structure via its catalytic function for the subsequent APE2 recruitment and activation for the continuation of SSB end resection (Lin et al., 2020). Interestingly, our observations (*Figure 1*) suggest that APE1 plays a downstream role in the ATR-Chk1 DDR, which is the recruitment of ATRIP onto ssDNA. To further support this distinct role of APE1, we have demonstrated evidence that the APE1-ssDNA and APE1-ATRIP interactions are important for ATRIP recruitment onto ssDNA *in vitro* (*Figure 3C*), and that ATRIP-interaction deficient mutant APE1 fails to recruit ATRIP onto ssDNA for subsequent ATR-Chk1 activation in *Xenopus* egg extracts (*Figure 3D*). Notably, the nuclease deficient ED APE1 is still proficient for ATRIP recruitment and ATR DDR in *Xenopus* egg extracts, similar to WT APE1 (*Figure 3D*). Together, these evidences support a non-catalytic role of APE1 via the direct recruitment of ATRIP onto ssDNA in the ATR-Chk1 DDR pathway.

A previous study has demonstrated that APE1 can incise the AP site within ssDNA in a sequence- and secondary-structure-dependent manner (Fan et al., 2006). Although this finding implies that that APE1 may associate with ssDNA, the molecular determinants of APE1 for ssDNA interaction is elusive. Our data in this study indicate that APE1 directly associates with ssDNA (>30nt length) and that the N-terminal 34 amino acids within APE1 are important for such direct ssDNA association (*Figure 2*). Although it remains unknown how the N-terminal domain of APE1 contributes to its 3’-5’ exonuclease activity but not AP endonuclease activity (*Figure 2-figure supplement 2C-D*), this is reminiscent of the regulation of APE2 3’-5’ exonuclease by its C-terminal Zf-GRF interaction with ssDNA (Wallace et al., 2017). It was previously reported that APE1 N-terminal domain associates with ssRNA to participate in RNA metabolism (Fantini et al., 2010). Future studies are needed to distinguish whether APE1 associates with ssDNA and ssRNA using similar or distinct mechanisms.

To the best of our knowledge, it is the first time showing that APE1 direclty associates with RPA via its two distinct motifs (*Figure 4*). A previous study mentioned that His-tagged human APE1 protein did not associate with purified untagged RPA protein using Ni-NTA-bead-based pulldown assays *in vitro* (Fan et al., 2006). Although we can’t explain the discrepancy with our result, we speculate that this may be due to different pullown methods and optimized experimental conditions. The two RPA bindings motifs within APE1 bind to RPA70 and RPA32, which is similar to the recently characterzied ETAA-RPA interaction (Bass et al., 2016; Haahr et al., 2016; Lee et al., 2016). Although RPA interaction is needed for the recruitment of ATR activator proteins ETAA1 and TopBP1 to damage sites and ssDNA for ATR activation (Acevedo et al., 2016; Bass et al., 2016; Haahr et al., 2016), our findings suggest that APE1 can associate with ssDNA in an RPA-independent manner, despite the APE1-RPA interaction. Future studies are warranted to find out the biological significance of the APE1-RPA interaction.

The data in this study support that APE1 directly associates and recruits ATRIP onto ssDNA independent of RPA in *in vitro* reconstitution systems, in addition to the well-established RPA-mediated recruitment of ATRIP onto ssDNA (Zou and Elledge, 2003) (*Figure 5A*). Furthermore, the RPA-mediated and APE1-mediated ATRIP recruitment onto ssDNA is neither dependent on nor exclusive to each other. However, in the *Xenopus* HSS system in which all RPA-interaction proteins are present (*Figure 5B*), the ATRIP recruitment onto RPA-coated ssDNA requires the presence of APE1 and its ATRIP-interaction binding N-terminal domain. We hypothesize that an unknown Protein X may negatively regulate ATRIP binding to RPA complex especially RPA70 N-terminal domain, which has been involved in several proteins’ recruitment such as ETAA1, Mre11, Nbs1, and Rad9. The Protein X’s negative regulation of ATRIP interaction may be counteracted by APE1 in the HSS system. Future studies will test this hypothesis. Taken together, we propose APE1 as a critical recruiter of ATRIP onto ssDNA in an RPA-independent manner to promote the ATR-Chk1 DDR pathway.

**Figure 5.**
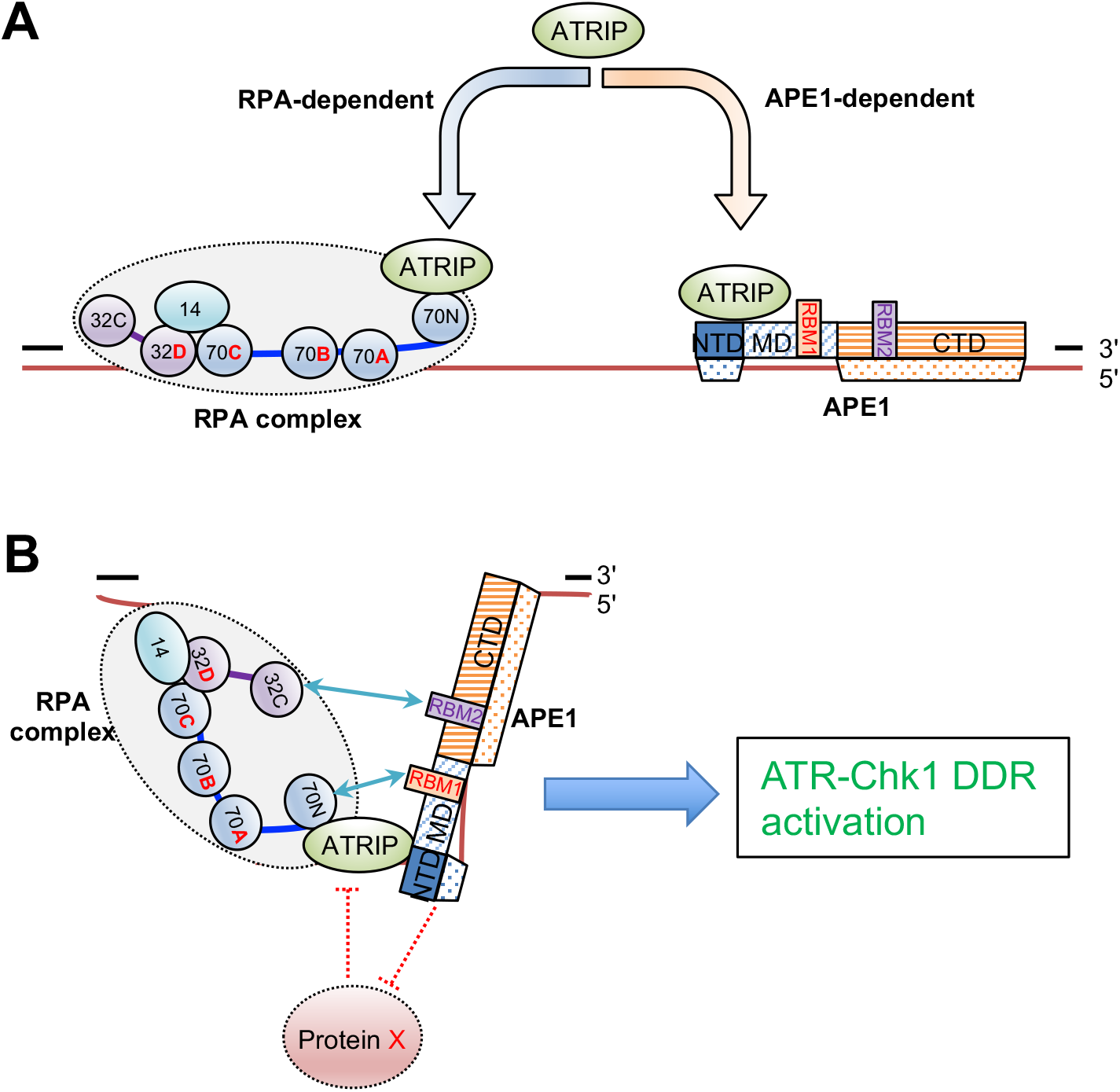
A working model of the distinct mechanism of how APE1 directly interacts and recruits ATRIP onto ssDNA independently *in vitro* and in a concerted fashion in Xenopus egg extracts for ATR DDR pathway. (**A**) RPA can recruit ATRIP to ssDNA in vitro. APE1 can substate RPA to promote ATRIP’s binding to ssDNA regions, which is dependent on APE1’s direct interaction with ssDNA and ATRIP. (**B**) APE1 is required in the recruitment of ATRIP to ssDNA in the ATR DDR in Xenopus egg extracts. APE1 interacts with RPA via two distinct motifs; however, APE1 interaction with ATRIP, but not its interaction with RPA nor APE1 nuclease, is important to recruit ATRIP onto ssDNA in Xenopus egg extracts to promote the ATR DDR activation.

## Materials and methods

### Key resources table

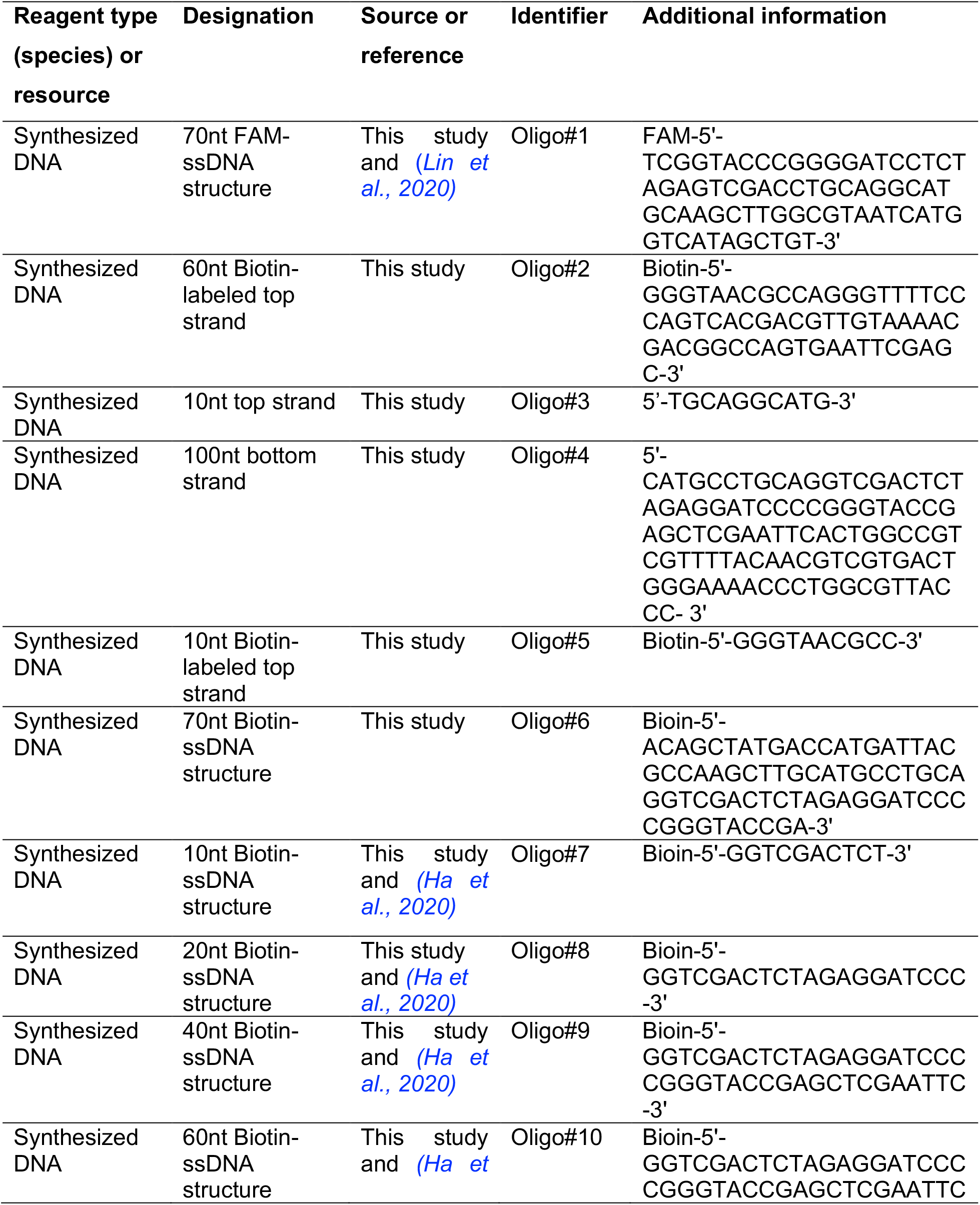
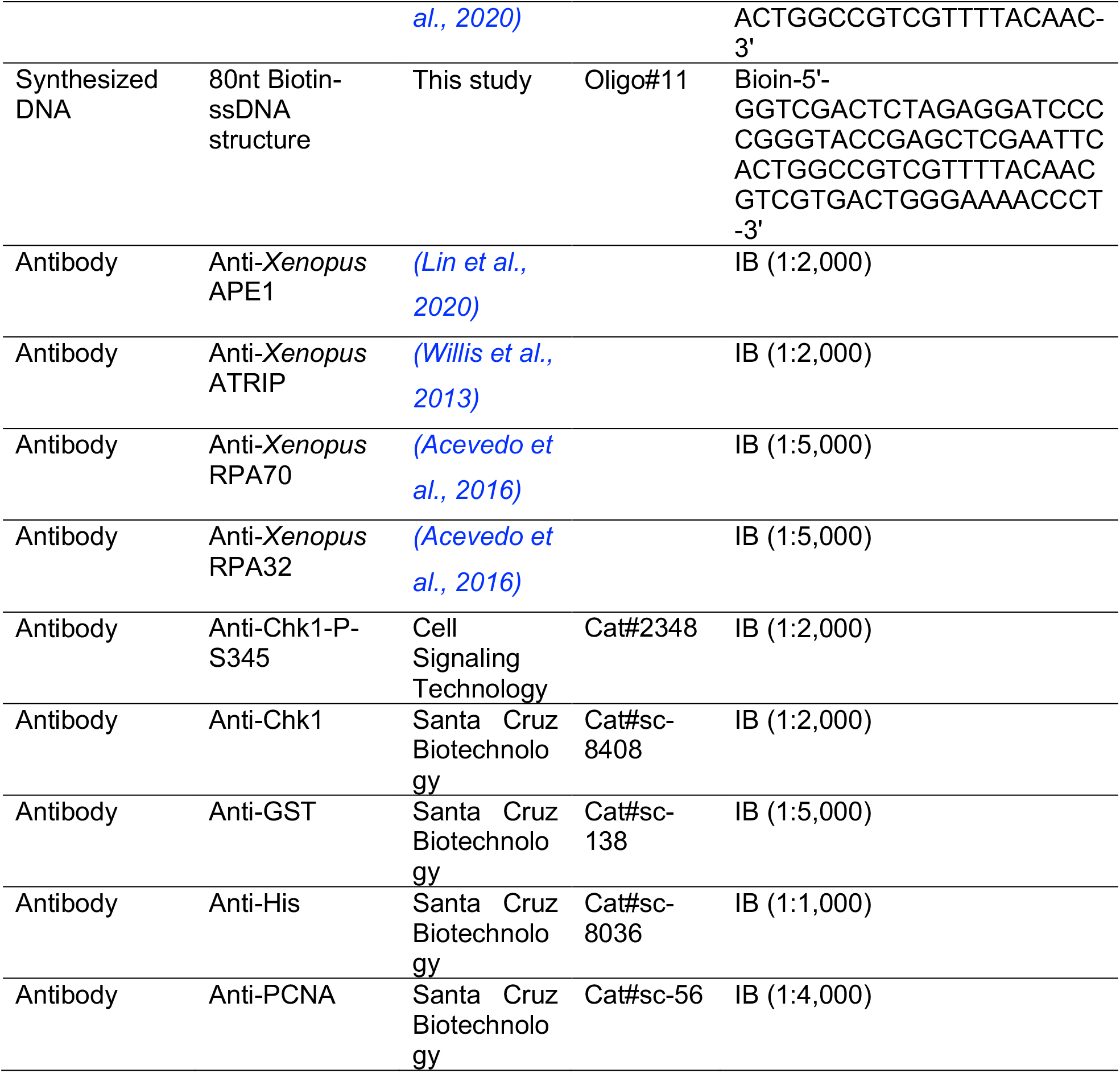

### Experimental procedures for *Xenopus* egg extracts, DDR signaling technology, and plasmid DNA bound fraction isolation in *Xenopus* egg extracts

The preparation of *Xenopus* HSS and immunodepletion of target proteins in HSS were described previously (Cupello et al., 2019; Cupello et al., 2016; Lebofsky et al., 2009; Lin et al., 2020; Willis et al., 2012). For the DDR signaling experiments, typically different plasmid DNA was mixed with HSS to final concentrations (e.g., 75ng/µL) for a 45-min incubation at room temperature using a protocol similar to previously described (Lin et al., 2019). Reaction mixture was added with sample buffer followed by examination via immunoblotting analysis. For DNA-bound protein isolation from HSS system, a detailed method has been described previously (Lin et al., 2018). Briefly, reaction mixture was diluted with egg lysis buffer followed by spinning through a sucrose cushion at 10,000 rpm at 4°C for 15 min. After aspiration of the supernatant, the DNA-bound protein factions were analyzed via immunoblotting analysis.

### Preparation of various plasmids and FAM/Biotin-labeled DNA structures

The preparation of control (CTL) plasmid and SSB plasmid was described previously (Lin et al., 2018; Lin et al., 2019). The SSB plasmid was treated by WT-GST-APE1 *in vitro* to create a gap plasmid structure (Lin et al., 2020).

The 39bp FAM-labeled dsDNA-AP structure for APE1 endonuclease assays was prepared as previously described (Lin et al., 2020). As shown previously (Lin et al., 2020), the 70bp FAM-dsDNA structure was prepared and treated with Nt.BstNBI and CIP to make the FAM-dsDNA-SSB for APE1 exonuclease assays. The FAM-dsDNA-SSB structure was purified from agarose via QIAquick gel extraction and then phenol–chloroform extraction. The 70nt FAM-ssDNA structure in *Figure 2E* was synthesize as Oligo#1.

The 100bp Biotin-dsDNA structure with a 30nt or 80nt ssDNA gap (30nt gap or 80nt gap) in the middle was created by annealing of three complementary oligos in same molar ratio at 95-100°C for 5 min followed by natural cooling down at room temperature for ∼30 min. For the 30nt ssDNA gap, the three complementary oligos are 60nt Biotin-labeled top strand Oligo#2, 10nt top stand Oligo#3, and 100nt bottom strand Oligo#4. For the 80nt ssDNA gap, the three complementary oligos are 10nt Biotin-labeled top strand Oligo#5, 10nt top stand Oligo#3, and 100nt bottom strand Oligo#4. The 70nt Biotin-ssDNA structure in *Figure 2D* and *Figure 2-figure supplement 1* was designed as Oligo#6. The Biotin-ssDNA structures with different lengths of ssDNA were synthesized as Oligo#7 (10nt), Oligo#8 (20nt), Oligo#9 (40nt), Oligo#10 (60nt), and Oligo#11 (80nt) and most of them were described previously (Ha et al., 2020).

### Recombinant DNA and proteins, and immunoblotting (IB) analysis

The preparation of recombinant WT, mutants (ED, DA, CA), and some fragments (AA35-316, AA101-316, AA101-200) of pGEX-4T1-APE1 was described previously (Lin et al., 2020). Other fragments of GST-APE1 (e.g., AA1-100, AA1-34, AA35-100) were generated by PCR of respective fragment and subcloned into pGEX-4T1. RBM1-M, RBM2-M, and RBM-DM pGEX-4T1-APE1 were mutated with QuikChange IIXL Site-Directed Mutagenesis kit (Agilent) and purified by QIAprep spin miniprep kit. His-RPA trimer expression plasmid was described previously (Acevedo et al., 2016). His-ATRIP expression plasmid was generated by PCR full length *Xenopus* ATRIP into pET28A at BamHI and XhoI sites. The induction/expression, purification and validation of GST or His-tagged recombinant proteins from BL21(DE3) E.coli cells (VWR Cat#80030-326) were performed following vendor’s standard protocol. Immunoblotting (IB) analysis was performed following similar methods described previously (Lin et al., 2020; Yan and Michael, 2009).

### GST pulldown assays and DNA binding assays

The GST-pull-down experiments were performed in an interaction buffer using similar methods as described previously (Lin et al., 2018; Lin et al., 2020). Methods for the DNA binding assays have been described previously (Lin et al., 2018; Lin et al., 2020). Briefly, Streptavidin Dynabeads coupled with various Biotin-labeled structures (e.g., Biotin-ssDNA, Biotin-dsDNA, or Biotin-dsDNA with ssDNA gap) was incubated with various recombinant proteins in a buffer (80 mM NaCl, 20 mM -glycerophosphate, 2.5 mM EGTA, 0.01% NP-40, 10 mM MgCl_2_, 100 ug/ml BSA, 10 mM DTT and 10 mM HEPES–KOH, pH 7.5) or with the HSS as indicated. After washing, the bead-bound fractions and Input samples were examined via immunoblotting analysis.

### *In vitro* endo/exonuclease assays

For *in vitro* APE1 endo/exonuclease assays, the FAM-dsDNA-AP structure or FAM-dsDNA-SSB structure was treated with WT, mutant or fragment of GST-APE1 in APE1 Reaction Buffer at 37°C, as described previously (Lin et al., 2020). The reactions were quenched with TBE-urea Sample Buffer and denatured for 5 min at 95°C. The samples were examined on TBE-urea PAGE gel and imaged with a Bio-Rad imager.

### Electrophoretic mobility shift assays (EMSA)

The EMSA assays for testing DNA-Protein interaction were similar to methods described previously (Lin et al., 2020). Briefly, an increasing concentration of proteins were incubated with 10 nM of FAM-labeled DNA structures in EMSA Reaction Buffer. Reactions were examined on a TBE native gel and imaged by a Bio-Rad imager.

### Material availability statement

Materials generated in this study can be accessed by contacting the corresponding author.

## Acknowledgments

We thank Drs. Matthew Michael, Karlene Cimprich, and Howard Lindsay for reagents. The Yan lab was supported, in part, by grants from the NIH/NCI (R01CA225637) and the NIH/NIEHS (R21ES032966), and funds from UNC Charlotte.

## Additional information

### Funding

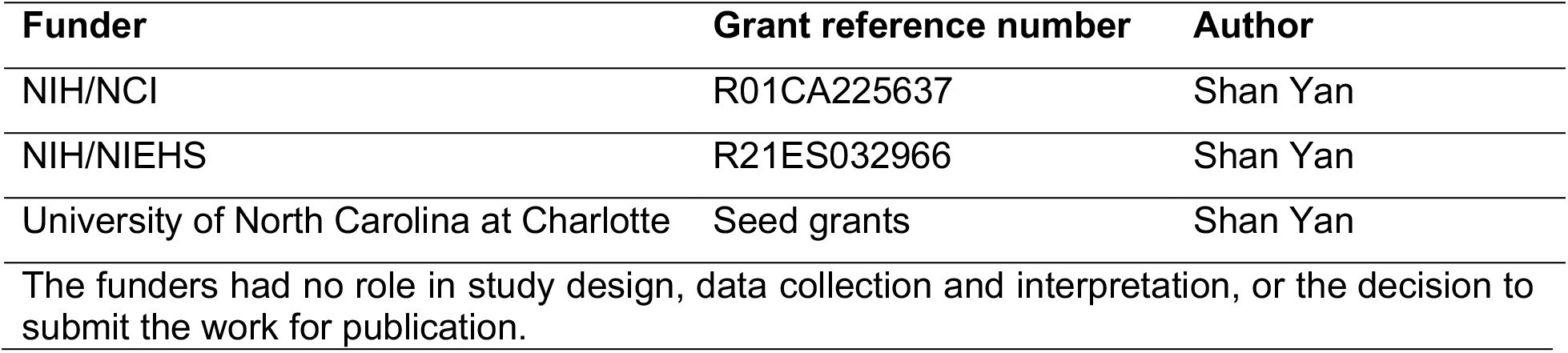

### Author contributions

Conceptualization, Y. Lin, S. Yan; data curation, Y. Lin, J. Li, H. Zhao, A. McMahon, S. Yan; funding acquisition, S. Yan; investigation, Y. Lin, J. Li, H. Zhao, A. McMahon, S. Yan; methodology, Y. Lin, S. Yan; supervision, S. Yan; writing-original draft, Y. Lin, S. Yan; writing-review and editing, Y. Lin, A. McMahon, S. Yan.

### Ethics

The care and use of X. laevis was approved by the Institutional Animal Care and Use Committee (IACUC) at University of North Carolina at Charlotte.

## Additional files

### Supplementary files

- MDAR checklist
- Source data 1. All original immunoblotting blotting images as well as IB-data with the uncropped images with the relevant bands clearly labelled and molecular weight markers.

### Data availability

All data generated or analyzed during this study are included in the manuscript and supporting files.

## Figures and Figure Legends

The online version of this article includes the following source data and figure supplements for Figure 1:

**Figure 1-figure supplement 1.**
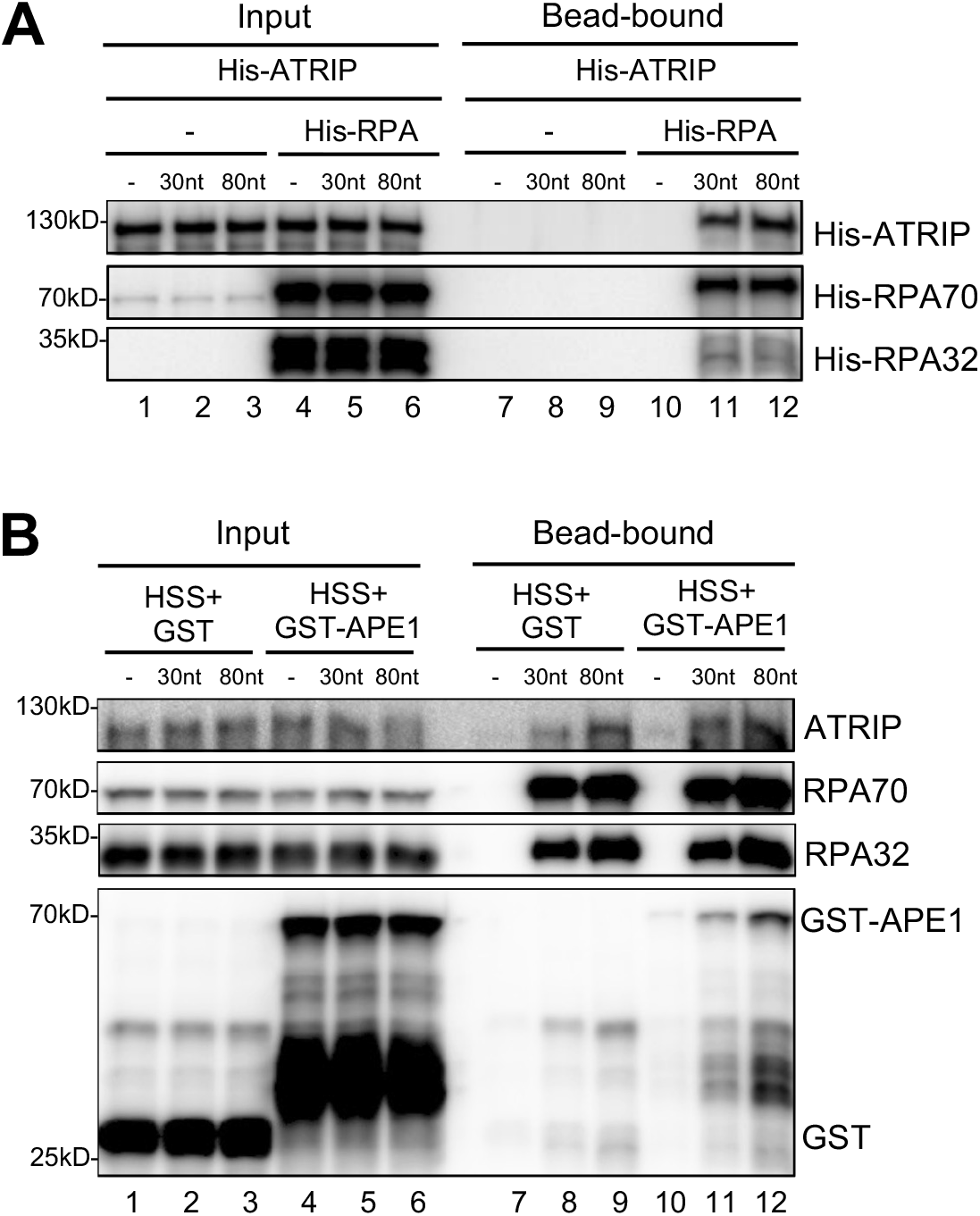
APE1 further promote ATRIP’s recruitment to RPA coated DNA. (**A**) Streptavidin beads coupled with same amount of Biotin-labeled dsDNA with ssDNA gap structures (30nt or 80nt) were added to an interaction buffer containing His-ATRIP, which was supplemented with/without His-RPA. DNA-bound fractions and total extract samples were examined via immunoblotting analysis as indicated. (**B**) Streptavidin beads coupled with equal moles of Biotin-labeled dsDNA with ssDNA gap structures (30nt or 80nt) were added to HSS, which was supplemented with GST or WT GST-APE1. DNA-bound fractions and total extract samples were examined via immunoblotting analysis as indicated.

The online version of this article includes the following source data and figure supplements for Figure 2:

**Figure 2-figure supplement 1.**
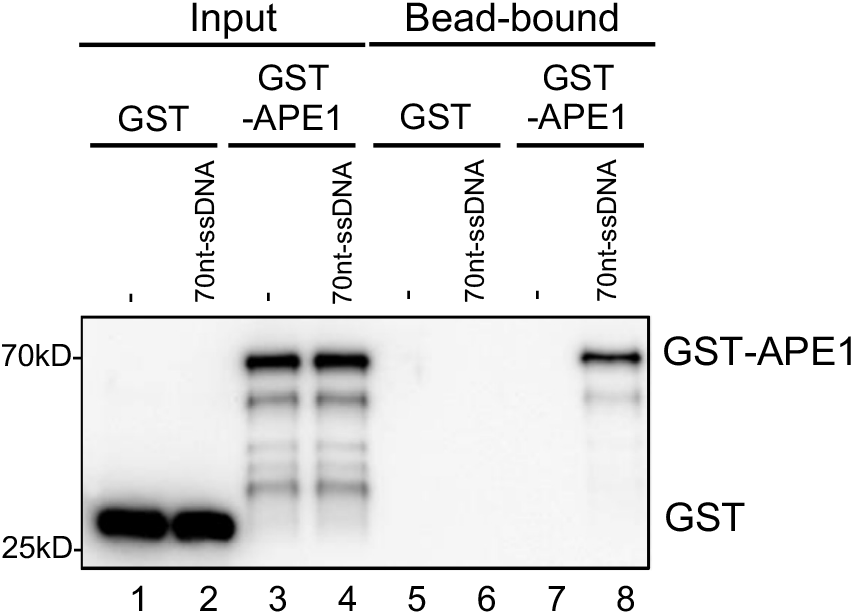
GST-APE1 but not GST associated with beads coupled with ssDNA. Streptavidin beads coupled with Biotin-labeled 70nt-ssDNA were added to interaction buffer, which was supplemented with GST or WT GST-APE1. DNA-bound fractions and Input samples were examined via immunoblotting analysis as indicated.

**Figure 2-figure supplement 2.**
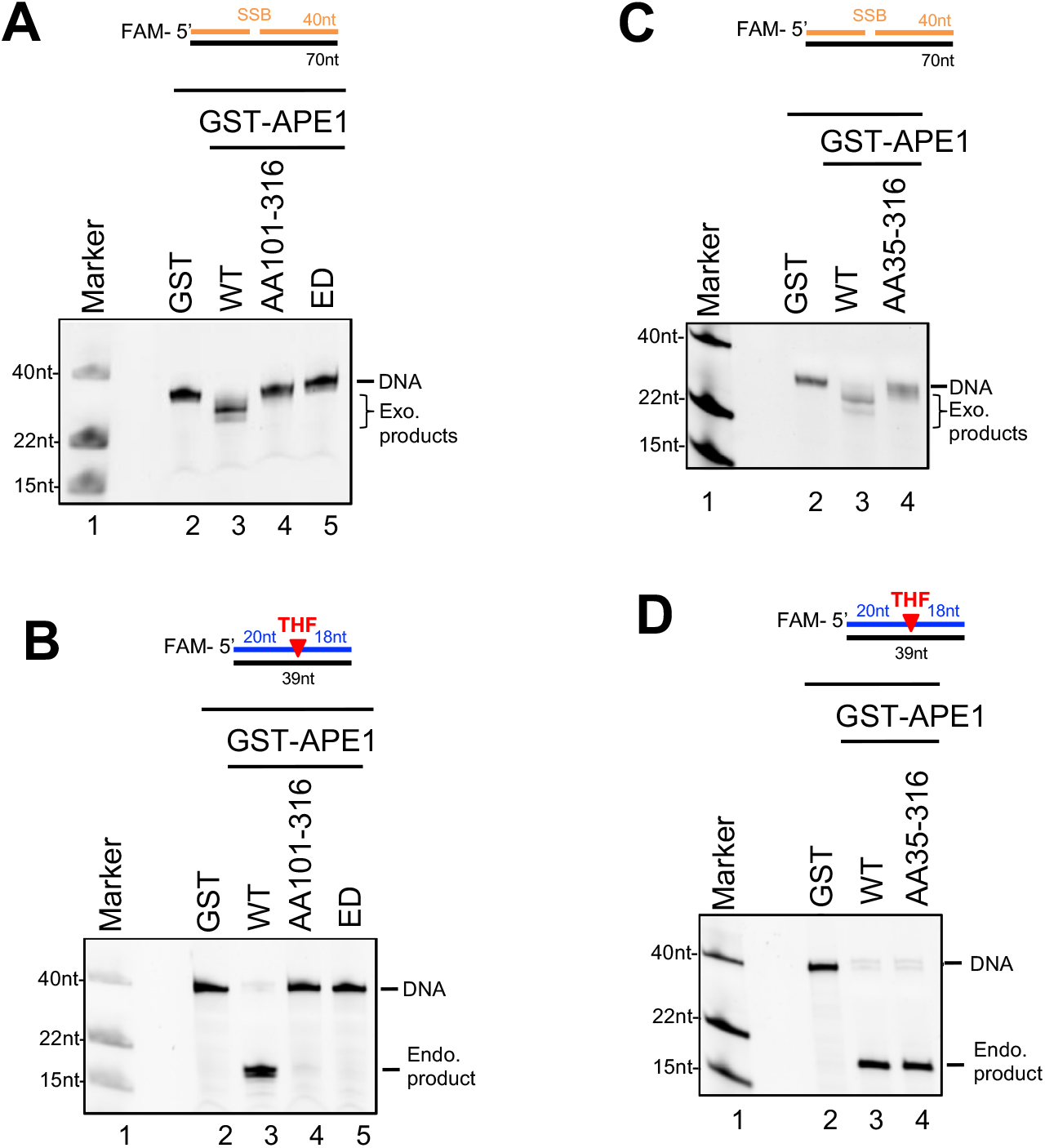
Endo/exonuclease activities of WT or various mutant/fragment of APE1 *in vitro*. (**A and C**) Various GST, or WT/mutant/fragment GST-APE1 was added to nuclease assay buffer containing the 70bp dsDNA-SSB structure for exonuclease activity assays. Samples were examined via denaturing urea PAGE electrophoresis and visualized. (**B and D**) Various GST, or WT/mutant/fragment GST-APE1 was added to nuclease assay buffer containing the 39bp dsDNA-AP structure for endonuclease activity assays. Samples were examined via denaturing urea PAGE electrophoresis and visualized.

The online version of this article includes the following source data and figure supplements for Figure 4:

**Figure 4-figure supplement 1.**
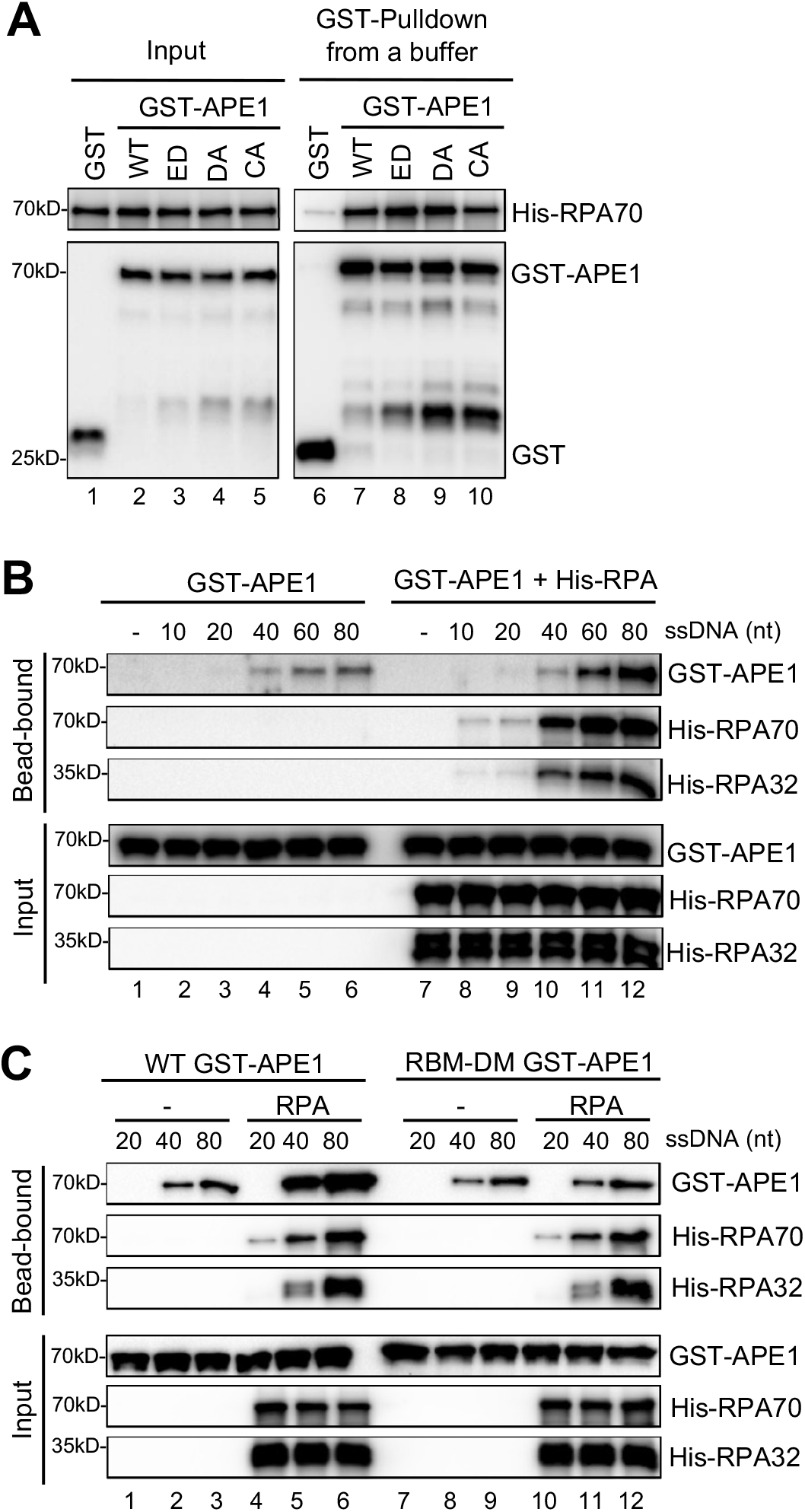
RPA-APE1 interaction promotes APE1 retention on ssDNA. (**A**) GST pulldown assays with GST, or WT/mutant GST-APE1 in an interaction buffer. The input and pulldown samples were examined via immunoblotting analysis. (**B**) Streptavidin beads coupled with Biotin-labeled ssDNA with different lengths were added to interaction buffer containing GST-APE1, which was supplemented with or without His-RPA protein. The input and pulldown samples were examined via immunoblotting analysis. (**C**) Streptavidin beads coupled with Biotin-labeled ssDNA with different lengths were added to interaction buffer containing WT/RBM-DM GST-APE1, which was supplemented with/without His-RPA protein complex. The input and pulldown samples were examined via immunoblotting analysis.

**Figure 4-figure supplement 2.**
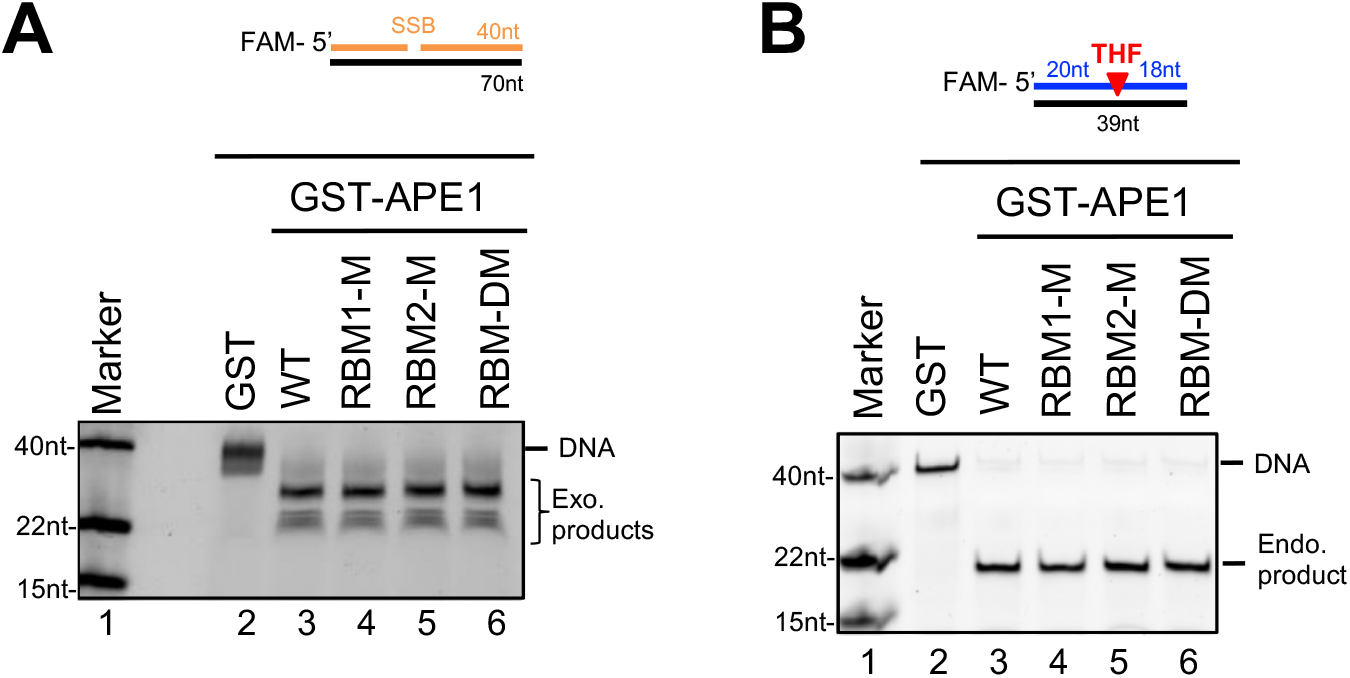
Endo/exonuclease activities of WT or various mutant/fragment of APE1 *in vitro*. (**A**) Various GST, or WT/mutant GST-APE1 was added to nuclease assay buffer containing the 70bp dsDNA-SSB structure for exonuclease activity assays. Samples were examined via denaturing urea PAGE electrophoresis and visualized. (**B**) Various GST, or WT/mutant GST-APE1 was added to nuclease assay buffer containing the 39bp dsDNA-AP structure for endonuclease activity assays. Samples were examined via denaturing urea PAGE electrophoresis and visualized.

